# Intestinal RICT-1/Rictor regulates larval germline progenitors via vitellogenins VIT-1 and VIT-3 in *C. elegans*

**DOI:** 10.1101/2025.01.08.632040

**Authors:** Anke Kloock, E. Jane Albert Hubbard

**Affiliations:** Department of Cell Biology, NYU Grossman School of Medicine, New York, NY 10016

**Keywords:** TORC2, Stem Cells, Rictor, Vitellogenin, germ line

## Abstract

Highly conserved signaling pathways communicate cellular and organismal nutrient conditions to proliferating pools of cells such as stem cells and tumors. One such pathway is the Target of Rapamycin (TOR) pathway. The TOR kinase exists in two complexes, TOR complex 1 (TORC1) and TOR complex 2 (TORC2). In the *Caenorhabditis elegans* hermaphrodite, TORC1 signaling cell-autonomously promotes the establishment of the larval germline Progenitor Zone (PZ). Here, we show that RICT-1, the sole *C. elegans* ortholog of the TORC2-specific component Rictor, also promotes expansion of the larval PZ through canonical TORC2 signaling. Unlike the TORC1 components, intestinal *rict-1* is both necessary and sufficient for full germline PZ pool establishment. Comparative RNA-seq on staged L4 larvae indicates that intestinal RICT-1 likely acts via vitellogenins. These intestinally produced yolk proteins provision oocytes, but they were not previously known to act in larval germ line development. Genetic analysis supports roles for *vit-1* and *vit-3* in a linear pathway with *rict-1*. Our results establish the *C. elegans* germ line as a fruitful model for investigating TORC2 and its connection to stem cells and lipid biology.

**Summary Statement:** RICT-1/Rictor in *C. elegans* promotes larval germline development via canonical TORC2 signaling, and intestinal RICT-1 acts via vitellogenins VIT-1 and VIT-3 to promote the establishment of the germline progenitor pool.

## INTRODUCTION

Highly conserved signaling pathways govern cellular responses to nutrients and metabolites. These responses are required for the proper regulation of pools of proliferating cells such as stem cells and tumor cells (Puca et al., 2022; Tu et al., 2023).

TOR is a highly conserved serine/threonine kinase that is conserved from yeast to mammals and integrates nutritional signals to regulate cellular responses (reviewed in (Liu and Sabatini, 2020)). TOR acts in two complexes bearing key subunits Raptor in TORC1 and Rictor in TORC2. Both complexes respond to nutritional signals and regulate many nutrient-dependent aspects of stem cell biology (reviewed in (Russell et al., 2011; Saba et al., 2021)). Albeit differently, both complexes respond to amino acid availability and impact lipid metabolism (reviewed in (Caron et al., 2015; Szwed et al., 2021; Zoncu et al., 2011)) and homeostasis (Kim and Chen, 2004). TORC2 furthermore promotes *de novo* fatty acid and lipid synthesis in mice (Guri et al., 2017) and worms (Jones et al., 2009; Soukas et al., 2009). In *C. elegans,* our prior work established that TORC1 components and/or downstream effectors affect germline stem cell proliferation and differentiation germ line-autonomously and in response to amino acid provision (Korta et al., 2012; Roy et al., 2018).

Reducing Rictor activity in rodents, worms, or flies results in small animals that develop slower (Guertin et al., 2006; Hietakangas and Cohen, 2007; Jones et al., 2009; Soukas et al., 2009). Rictor mutants in rodents and worms accumulate fat (Frei et al., 2023; Jones et al., 2009; Soukas et al., 2009; Yen et al., 2010), suggesting Rictor may facilitate trafficking of lipids from the intestine to other organs (Jones et al., 2009; Soukas et al., 2009). Adult *C. elegans* hermaphrodites bearing mutations in *rict-1*, the sole Rictor ortholog (Jones et al., 2009), exhibit reduced intestinal expression of the vitellogenin gene *vit-3* (Breen et al., 2024; Dowen et al., 2016) and produce fewer offspring (Jones et al., 2009; Soukas et al., 2009; Webster et al., 2013).

A pool of stem and progenitor cells, known collectively as the progenitor zone (PZ) pool, exists in the distal part of the hermaphrodite gonad (reviewed in (Hubbard and Schedl, 2019)), and the PZ pool is related to offspring production in sperm-replete worms (Agarwal et al., 2018; Kocsisova et al., 2019). The PZ is maintained by Notch pathway signaling via direct activation of downstream targets *sygl-1* and *lst-1* (Chen et al., 2020; Kershner et al., 2014) that interfere with three partially redundant differentiation pathways (Mohammad et al., 2018).

Our results show that *rict-1* mutant worms accumulate half the normal number of cells in the PZ during larval stages and have a reduced number of offspring. As in other phenotypic contexts, *rict-1* acts via *sinh-1* and *sgk-1*. However, in this role, *rict-1* does not act via the IIS or TGF-ß pathways which are implicated in larval PZ expansion (Dalfó et al., 2012; Michaelson et al., 2010; Pekar et al., 2017) and in *rict-1* control of dauer entry (O’Donnell et al., 2018). Loss of *rict-1* reduces the average mitotic index in larvae and likely acts in parallel to GLP-1. We show that intestinal RICT-1 is sufficient and necessary for full establishment of the adult PZ. Our transcriptomics analysis indicates that during the L4 larval stage, *rict-1* regulates expression of several members of the conserved ApolipoproteinB-like vitellogenin family (reviewed in (Perez and Lehner, 2019)), and that *vit-1* and *vit-3* contribute to expansion of the progenitor pool, likely downstream of *rict-1*.

## RESULTS

### RICT-1 is required for normal germline development

To explore effects of TORC2 on the germ line, we examined the effects of reduced *rict-1.* We first assessed phenotypes in two *rict-1* mutants (*ft7* in the third exon and *mg360* in the 11^th^ exon (Jones et al., 2009; Soukas et al., 2009)), in 6 additional alleles that we generated using CRISPR-Cas9 genome editing, and in an additional deletion mutant (Breen et al., 2024). Among three different isoforms *ft7* only affects isoforms A and B, while *mg360* affects all three (Fig. 1A). We generated four CRISPR mutants with Stop-In cassettes (Wang et al., 2018) inserted in the regions of each of the SNPs: *na112* and *na113* in the third exon, and *na11*4 and *na115* in the 11^th^ exon. We also generated two additional mutants: one that deletes 241bp of the 5’UTR and 29bp of the first exon (*na116*), and one that deletes 520bp in total across the first three exons (*na119*). We used the Stop-In strategy so as not to interfere with the *pqn-32* locus that resides within the intron between exons 10 and 11. We furthermore assessed a mutant that deletes both *rict-1* and *pqn-32* (Breen et al., 2024).

**Figure 1:**
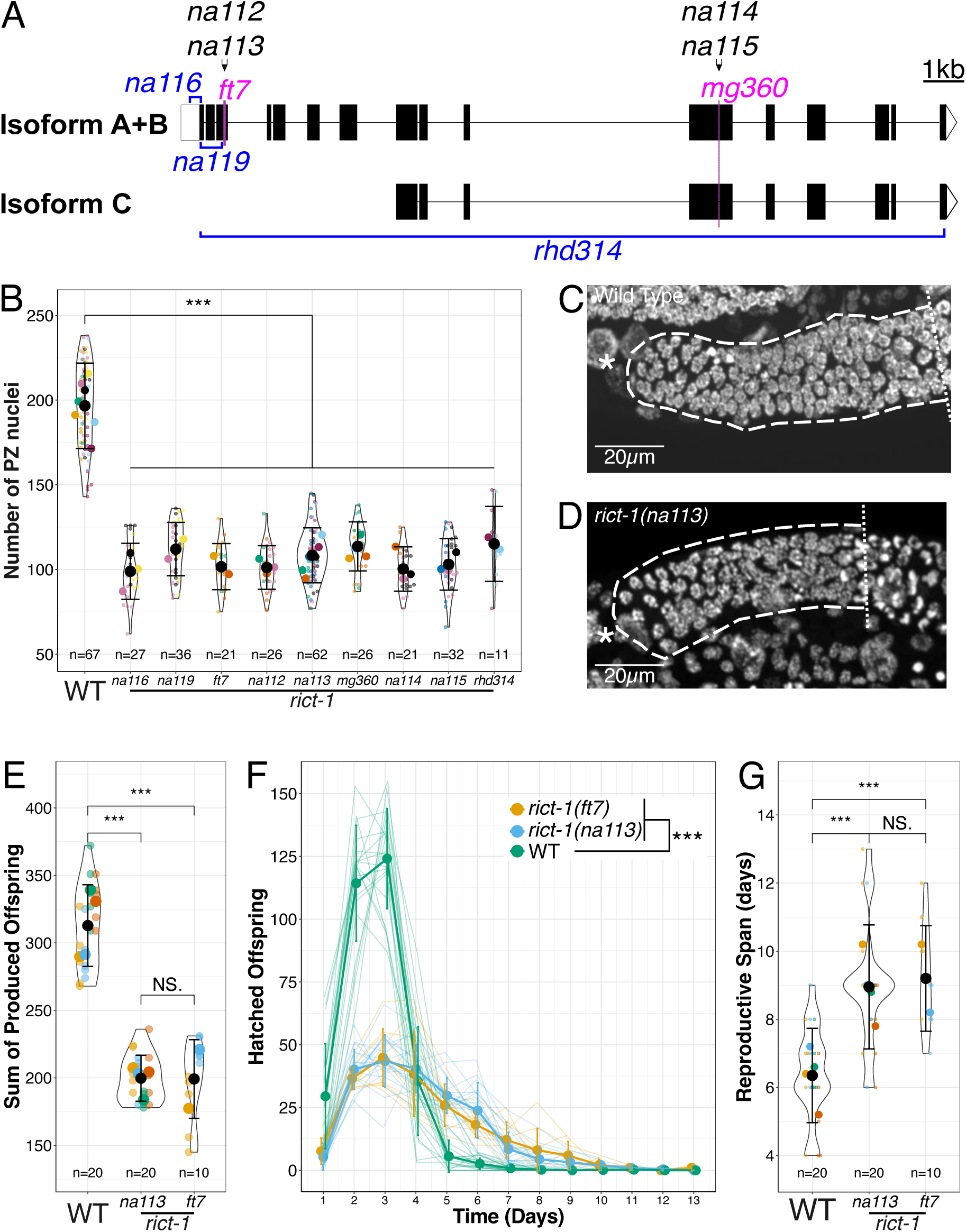
*rict-1* regulates reproduction and the germline progenitor zone. (A) Overview of the nine different *rict-1* mutants used in this study (two SNP mutations in magenta, four Stop-in mutants in black, and three knock-out mutants in blue). Scale bar indicates 1kb. Alleles *mg360*, *na114*, *na115,* and *rhd314* affect all isoforms. (B) Number of PZ nuclei for wild type and nine tested *rict-1* mutants. (C, D) Micrographs of representative wild type (C) and *rict-1(na113)* mutant (D) progenitor zones. (E) Average of hatched offspring per adult over a lifetime. (F) Hatched offspring per adult over time; the thick lines indicate the average per strain, while the thinner lines indicate single worms. Error bars indicate standard error of the mean. (G) Reproductive period in days. (B, E, G) Black circles indicate the means across independent replicate experiments (these replicate experiments are shown in different colors), while the smallest circles indicate gonad arms (n = number of gonad arms scored), also summarized below each violin. The violin displays the overall distribution of data points. Error bars indicate the standard error of the mean. Statistical results and p-values are summarized in Tables S6 and S7, respectively. The asterisks indicate *** p<0.001, ** p<0.01, * p<0.05. Panel B, E, and G were analyzed with a Linear Mixed Effects Model, Panel F was analyzed with a Generalized Linear Model, followed up with a Tukey Post Hoc test where appropriate.

To gauge the effects of *rict-1* on larval accumulation of the stem/progenitor pool, we counted the number of nuclei per distal gonad arm in the distal progenitor zone (PZ) at the L4-to-adult molt. We chose to assess the progenitor pool at the molt because this represents a developmental timepoint that is independent of growth rate and is at the start of the early adult steady-state PZ pool (reviewed in (Hubbard and Schedl, 2019)). All nine *rict-1* mutants displayed a similar significant reduction in the number of PZ nuclei, reaching half of the wild-type level (average 106.2 ± 1.00 across all nine alleles versus 196.6 ± 3.08 in the wild type) (Fig. 1B-D). Given that all *rict-1* mutants behaved similarly, our subsequent studies used the *na113* allele, indicated as *rict-1(-)* hereafter.

The brood size in mated worms is correlates with the number of PZ nuclei (Agarwal et al., 2018; Kocsisova et al., 2019). To determine whether the PZ pool reduction we observed in the *rict-1* mutants is associated with reproductive defects, we assessed live broods and, consistent with previous results (Jones et al., 2009; Soukas et al., 2009), we observed a significantly smaller number of hatched offspring for both *rict-1* alleles tested (199.2 ± 9.21 for *rict-1(ft7)* and 199.8 ± 3.81 for *rict-1(na113)*) compared with the wild type (312.8 ± 6.77) (Fig. 1E,F). We also observed that *rict-1* mutants display an extended reproductive span of ∼9 days compared with ∼6 days in the wild type (9.0 ± 0.41 days for *rict-1(na113)*, 9.2 ± 0.49 for *rict-1(ft7)*, and 6.4 ± 0.31 for the wild type (Fig. 1F,G)), consistent with previous results (Webster et al., 2013). Since hermaphrodite brood size is sperm limited (Hodgkin and Barnes, 1991), and the correlation with the PZ can only be assessed in the presence of replete sperm, we assessed live broods in mated worms. We found that, unlike the wild type, where male sperm provision significantly increased offspring number, the number of hatched offspring is not increased in mated *rict-1* mutants (Fig. S1). Despite the extended reproductive period, the overall sum of hatched brood of *rict-1* mutants is still reduced relative to wild type (Fig. 1F). We conclude that RICT-1 promotes normal germline development, and we focused on the PZ pool phenotype as this is a primary early aspect of germline development.

### RICT-1 acts through canonical TORC2 to establish the early adult PZ pool

We next assessed known pathway components or downstream effectors of the canonical TORC2 complex. Canonically, Rictor and SINH-1 are in the TORC2 complex, and TORC2 activates SGK (reviewed in (Blackwell et al., 2019; Jang et al., 2022)).

In *C. elegans*, SINH-1 acts similarly to Rictor in many phenotypic contexts (Aspernig et al., 2019; Emans et al., 2023; McLachlan et al., 2022). To test if RICT-1 is acting through canonical TORC2 to establish the larval PZ, we assessed the number of PZ nuclei in worms bearing mutations in *rict-1(-)* or *sinh-1(-)* or both (Fig. 2A). We found that the number of PZ nuclei was similarly reduced in each single mutant and in the double mutant (104.7 ± 3.35, 114.3 ± 3.68, and 107.8 ± 3.25, respectively). These results are consistent with RICT-1 and SINH-1 acting in the same complex.

**Figure 2:**
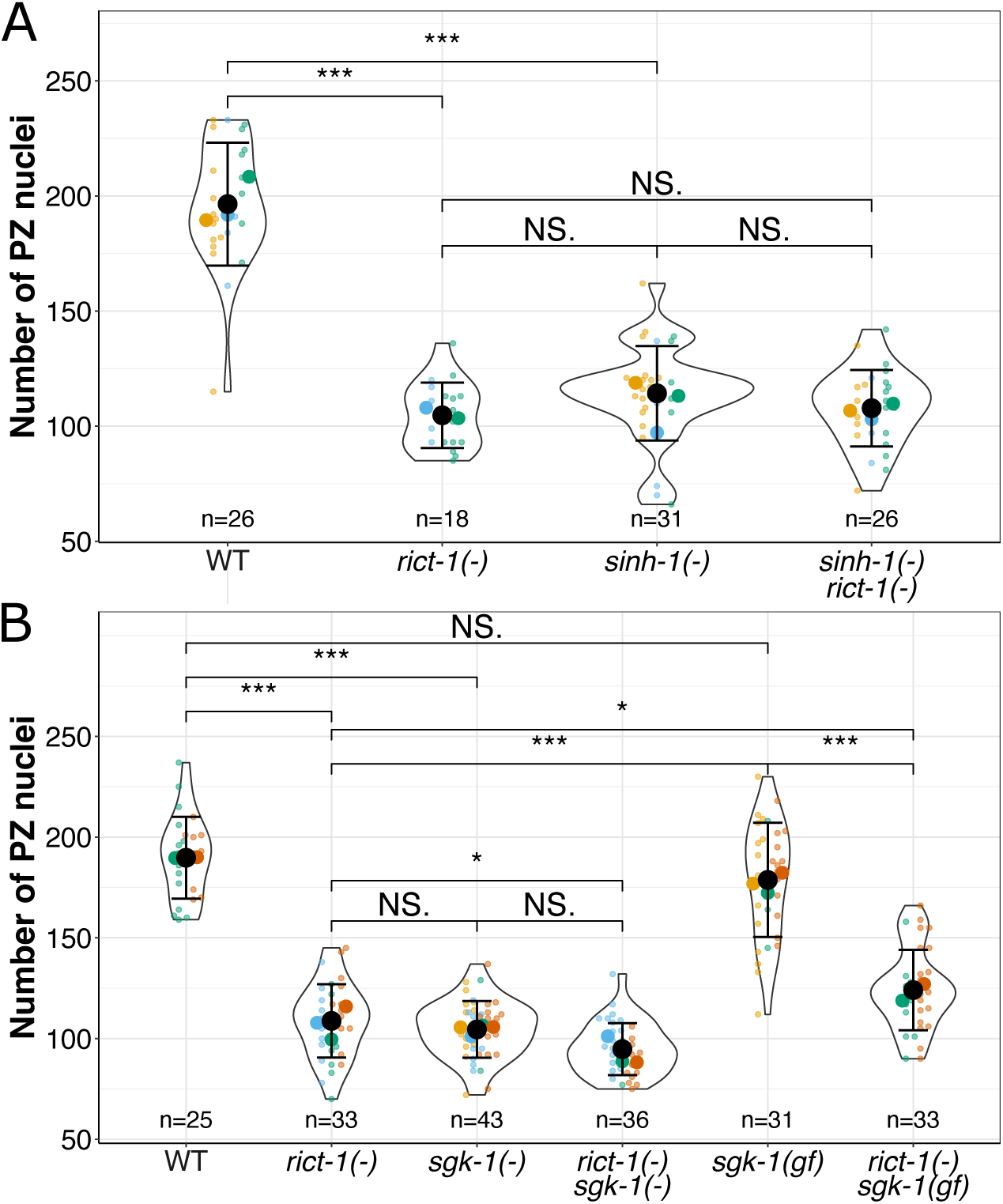
RICT-1 likely acts via canonical TORC2 signaling pathway. (A, B) Number of PZ nuclei in the wild type (WT) and indicated mutant genotypes. Black circles indicate the means across independent replicate experiments (these replicate experiments are shown in different colors), while the smallest circles indicate gonad arms (n = number of gonad arms scored), also summarized below each violin. The violin displays the overall distribution of data points. Error bars indicate the standard error of the mean. Statistical results and p-values are summarized in Tables S6 and S7, respectively. The asterisks indicate *** p<0.001, ** p<0.01, * p<0.05. The asterisks indicate *** p<0.001, ** p<0.01, * p<0.05 of a Linear Mixed Effects Model with a Tukey Post Hoc Test.

In *C. elegans*, SGK-1 acts genetically downstream of Rictor in many phenotypic contexts (Aspernig et al., 2019; Dowen et al., 2016; Emans et al., 2023; Gatsi et al., 2014; Jones et al., 2009; Ruf et al., 2013; Soukas et al., 2009; Webster et al., 2013). To determine whether a similar genetic interaction exists for PZ establishment, we compared PZ counts in the null single and double mutants. We found that the number of PZ nuclei is similar in *rict-1* or *sgk-1* single mutants (108.8 ± 3.17 and 104.5 ± 2.15 for *rict-1(-)* and *sgk-1(-)*, respectively) and in the *rict-1; sgk-1* double mutant (94.7 ± 2.15) (Fig. 2B). While PZ counts in each single mutant do not differ, the *rict-1*; *sgk-1* double mutant is modestly, but significantly, lower than in the *rict-1* mutant alone (p=0.024). This result suggests that while *sgk-1* is likely acting downstream of *rict-1*, it may have additional *rict-1*-independent minor effects on the PZ phenotype independent of *rict-1* (Fig. 2B).

An *sgk-1* gain-of-function allele (*gf*) (Jones et al., 2009) fully (Jones et al., 2009; Sakai et al., 2017) or partially (Aspernig et al., 2019; Ruf et al., 2013) suppresses *rict-1* mutant phenotypes, consistent with a role for SGK-1 downstream of RICT-1 (Jones et al., 2009; Sakai et al., 2017). To determine whether the *sgk-1(gf)* mutant suppresses the *rict-1* mutant PZ phenotype, we examined *sgk-1(gf)* alone and in combination with *rict-1(-).* We found that *sgk-1(gf)* alone displays a PZ count similar to the wild type (178.7 ± 5.10 versus 189.8 ± 4.06). As expected if SGK-1 were acting downstream of RICT-1, we found significant, though partial, suppression of the *rict-1* PZ phenotype (to 124.1 ± 3.48 PZ nuclei). Taking the loss- and gain-of-function results together, we conclude that SGK-1 likely acts downstream of RICT-1, even though the two may have other minor independent outputs that affect the PZ phenotype.

### RICT-1 does not act via DAF-2/IIS or DAF-7/TGF-ß pathways to promote accumulation of the larval progenitor pool

In the phenotypic context of dauer entry, intestinal *rict-1* signals via neuronal DAF-28 Insulin and DAF-7 TGF-ß from ASI and ASJ (O’Donnell et al., 2018). This finding was of particular interest to us since we previously characterized roles for both the IIS and TGF-ß pathways in establishing the PZ pool (Dalfó et al., 2012; Michaelson et al., 2010; Pekar et al., 2017). In both pathways, these same neurons are implicated: *ins-3* is implicated upstream of DAF-2 for PZ phenotypes and is expressed in ASI and ASJ (Ghaddar et al., 2023; Michaelson et al., 2010), while ASI is implicated by cell ablation for the DAF-7 pathway regulation of the germ line stem cell niche (Dalfó et al., 2012). Importantly, the PZ accumulation defect upon reduced *daf-2* or *daf-7* is fully suppressible by loss of *daf-16* or *daf-3*, respectively.

If the relevant IIS and/or TGF-ß pathways mediate the effects of *rict-1* on the PZ, we would expect loss of *daf-16* and/or *daf-3* to suppress the *rict-1* mutant PZ defect. To test this hypothesis, we constructed strains bearing double or triple mutants of *rict-1(-)* with *daf-16(-)* and/or *daf-3(-)* and counted the number of nuclei in the PZ pool. We found that neither *daf-16(-)* nor *daf-3(-)* alone strongly suppressed the PZ defect in the *rict-1* mutant (Fig. 3A). The PZ of the *daf-16(-); rict-1(-)* averaged 119.1 ± 2.91 (compared with 97.6 ± 2.73 for *rict-1(-)* alone). While this small difference is statistically significant, we note that *daf-16(-)* has a greater (though not statistically significant) average number of PZ nuclei than the wild type (Fig. 3A). For the DAF-7/TGFß pathway, we found that the PZ pool of *daf-3(-); rict-1(-)* was not significantly different from the *rict-1* mutant alone. Most important, the triple mutant *daf-16(-); rict-1(-); daf-3(-)* averages the same number of PZ nuclei (109.0 ± 2.54) as either double mutant or the *rict-1(-)* mutant alone (Fig. 3A). We conclude from these results that neither DAF-2/IIS (via DAF-16) nor DAF-7/TGFß pathways mediate the effects of RICT-1 on PZ accumulation.

**Figure 3:**
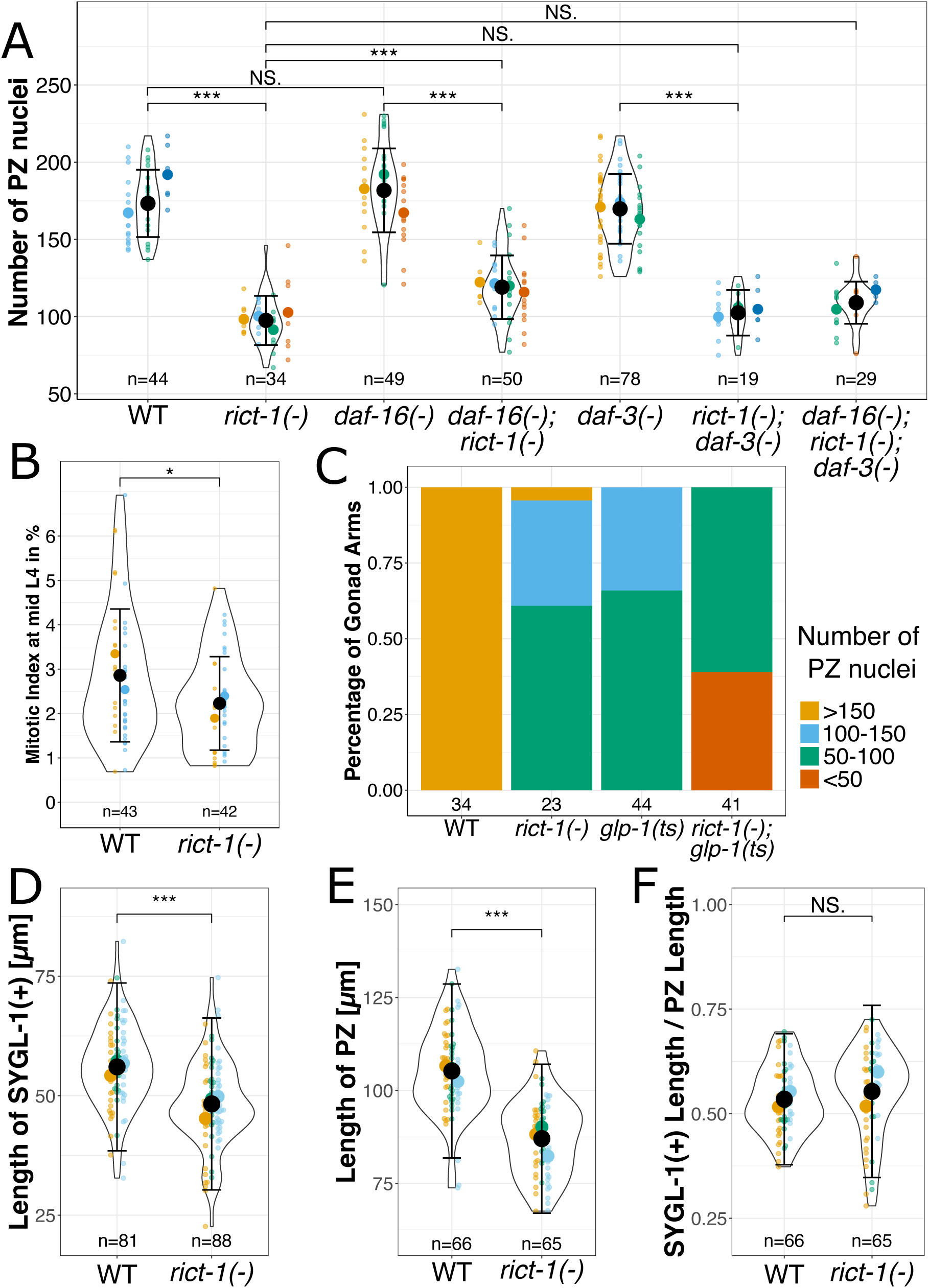
RICT-1 does not act via the IIS or the DAF-7/TGF-ß pathway but regulates larval cell cycle and not stem cell fate. (A) Number of PZ nuclei in the wild type (WT) and indicated mutants. (B) Mitotic index at the mid-L4 in the WT and *rict-1(-)* mutant. (C) The percentage of gonad arms displaying the indicated range in numbers of PZ nuclei. (D, E, F) The Length of the SYGL-1(+) (D) and PZ (E) zones, and their ratio (F) in WT and *rict-1(-)* mutants. (A, B, D-F) Black circles indicate the means across independent replicate experiments (these replicate experiments are shown in different colors), while the smallest circles indicate gonad arms (n = number of gonad arms scored), also summarized below each violin. The violin displays the overall distribution of data points. Error bars indicate the standard error of the mean. Statistical results and p-values are summarized in Tables S6 and S7, respectively. The asterisks indicate *** p<0.001, ** p<0.01, * p<0.05 of a Linear Mixed Effects Model with a Tukey Post Hoc Test. (C) Stacked bar plots represent the ratio of gonad arms that fall into different categories, indicated on the right (yellow for gonad arms with >150 PZ nuclei, light blue for gonad arms with 100-150 PZ nuclei, green for gonad arms with 50-100 PZ nuclei, and orange for gonad arms with <50 PZ nuclei. Individual examined gonad arms are indicated below each bar.

### RICT-1 promotes larval cell cycle progression, but does not enhance the Glp-1-like “loss of progenitor pool” phenotype

Our prior work established that *rsks-1/*S6K affects both the cell cycle and cell fate in the PZ (Korta et al., 2012; Roy et al., 2016). Loss of *rsks-1* reduces both larval mitotic index and S phase index, as well as on the proportion of cells in G1 or G2, suggesting a generally slower cell cycle in the mutant. These reductions were observed in the mid-L4 stage larvae when the PZ is still expanding, however no differences were observed in the adult homeostatic stage when the cell cycle is already slowed relative to larval stages. Moreover, in combination with a reduction of *glp-1* (using a temperature-sensitive allele at a semi-permissive temperature), loss of *rsks-1* had a dramatic effect on the PZ such that in over 50% of *glp-1(ts) rsks-1(sv31)* double mutants at 20°C, all germ cells had differentiated, mimicking a complete loss of *glp-1* and demonstrating a parallel role for *rsks-1* in cell fate (Korta et al., 2012; Roy et al., 2016).

To determine how *rict-1* affects cell cycle and cell fate, we assessed the mitotic index in the *rict-1(-)* mutant in mid-L4 stage larvae and found that, similar to *rsks-1,* the larval mitotic index was reduced in *rict-1(-)* mutants (2.23 ± 0.16 for *rict-1(-)* mutants versus 2.85 ± 0.23 for wild-type (Fig. 3B)). Similar to prior results with *rsks-1, rict-1* does not affect the mitotic index in adults (Fig. S2A).

Furthermore, we assessed the genetic interaction with *glp-1*. Unlike what was observed with *rsks-1*, the double *glp-1(ts); rict-1(-)* mutant did not cause differentiation of the entire PZ (Fig. 3C). However, the double mutant (*rict-1(-); glp-1(ts)*) showed a minor increase in the percentage of gonad arms with fewer than 50 PZ nuclei (39% of gonad arms for the *rict-1(-); glp-1(ts)* mutant with <50 PZ nuclei, versus 0% for each single mutant) (Fig. 3B). This suggests that RICT-1 acts in parallel to *glp-1* (Fig. 3B). Furthermore, the number of PZ nuclei for the double mutant of *rict-1(-)* and *glp-1(ts)* is additive (Fig. S2B). Together, these results support the conclusion that RICT-1 is acting to promote robust larval cell cycle progression but does not affect the stem cell fate decision.

Within the PZ pool there are distal SYGL-1(+) stem cells (Shin et al., 2017) and more proximal SYGL-1(-) progenitors that are completing a final round of mitosis (Fox and Schedl, 2015). We asked whether loss of *rict-1* affects these early adult pools independently. Using as a readout the distance from the distal end of the gonad to the proximal border of each pool, we found that *rict-1(-)* mutants had a significantly shorter SYGL-1(+) zone than wild-type worms (p<0.001, 48.3 ± 0.96 µm versus 56.0 ± 0.96 µm, respectively) (Fig. 3C). The length of the PZ was also reduced in *rict-1(-)* mutants in comparison to wild-type worms (p<0.001, 87.0 ± 1.24 versus 105.2 ± 1.44, respectively) (Fig. 3D). When we assessed the relative length of the SYGL-1(+) zone to the PZ, we observed that in both strains the SYGL-1(+) zone is about half the length of the PZ (0.55 ± 0.013 for *rict-1(-)* mutants and 0.53 ± 0.009 for the wild type) (Fig. 3E). These results suggest that the loss of robust cell cycle progression in larval stages affects both the stem and non-stem-progenitor portions of the adult PZ pool.

### Intestinal RICT-1 contributes to PZ pool establishment

In several different phenotypic contexts, *rict-1* acts from the intestine. These contexts include dauer entry and foraging behavior (O’Donnell et al., 2018), associative learning (Sakai et al., 2017), fat accumulation (Jones et al., 2009; Soukas et al., 2009), and food seeking (McLachlan et al., 2022). Single cell RNA-seq analyses show that *rict-1* mRNA is widely expressed but is most highly abundant in the intestine (Fig. S3A from (Cao et al., 2017; Hutter and Suh, 2016)).

We therefore asked whether intestinal RICT-1 might influence PZ pool establishment. To test this, we depleted RICT-1 in the intestine using tissue-specific auxin-mediated protein degradation(Zhang et al., 2015). We generated a sAID::RICT-1 fusion (*rict-1(na117)*) by CRISPR-Cas9 genome editing, and determined that it does not impact protein function, as the number of PZ nuclei is not significantly different from that of wild-type worms (Fig. S3B). We then crossed this allele into a strain in which TIR1 is driven by the intestine-specific promoter *ges-1p.* We observed that in the presence of auxin, the number of PZ nuclei is significantly reduced (153.3 ± 3.23 with auxin vs. 189.7 ± 3.21 without auxin). Neither the vehicle (EtOH) nor auxin alone affected the PZ counts of the other strains tested (wild type, *rict-1(-)*, *rict-1(na117*[sAID::RICT-1]) or *ieSi61*[*ges-1p*::TIR1] alone), indicating that the lower PZ count we observed in the presence of the TIR1 and auxin is due to auxin-induced protein degradation and not to auxin-induced stress (Fig. 4A).

**Figure 4:**
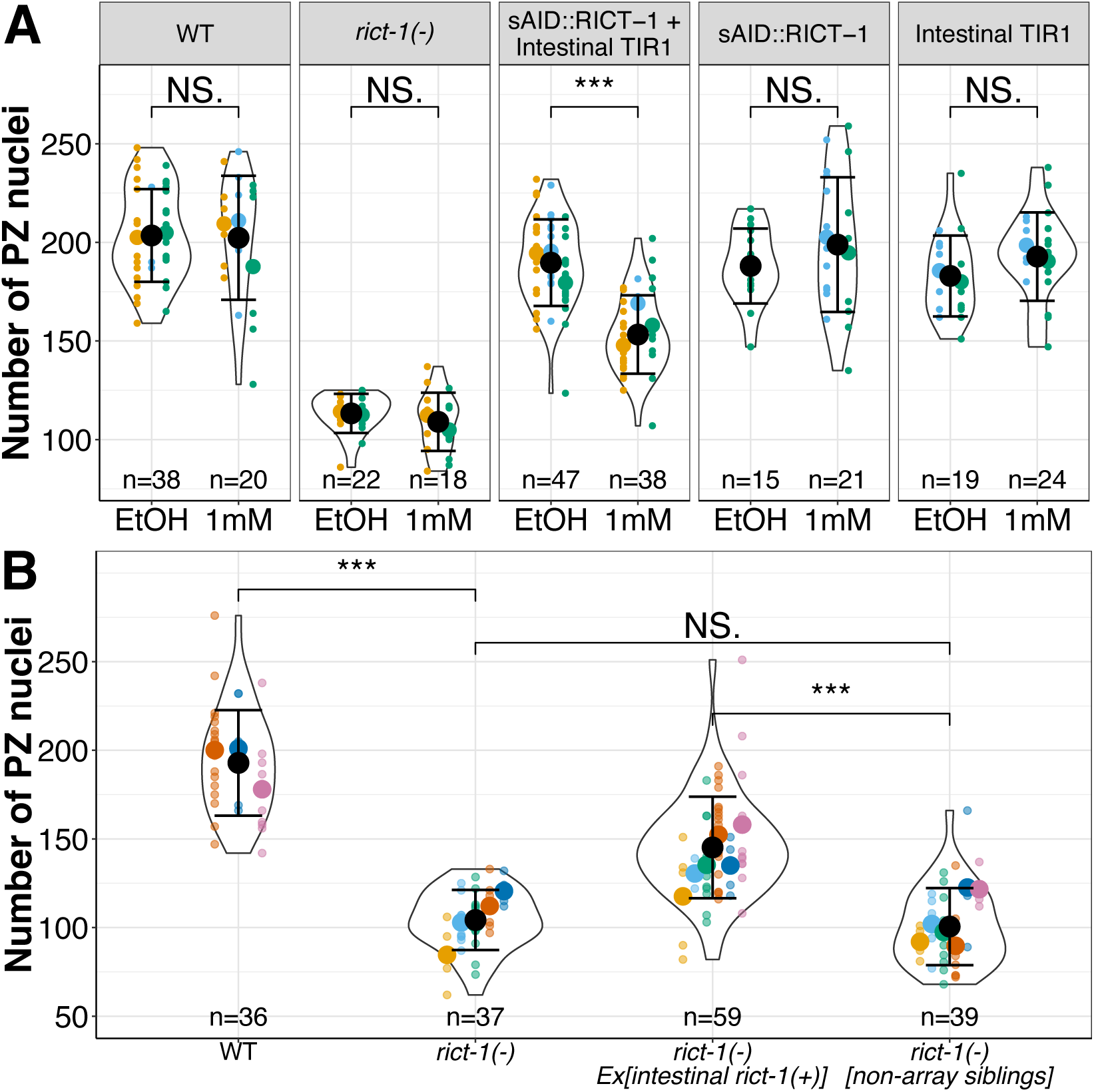
Intestinal RICT-1 is necessary and sufficient for full establishment of the PZ. (A) Numbers of PZ nuclei in the presence of the vehicle (EtOH) and auxin in the wild type (WT) and *rict-1(-)* and in strains bearing sAID::RICT-1 and/or the intestinal TIR1 (*ieSi61*). (B) Numbers of PZ nuclei in the wild type (WT) and in array bearing and non-array bearing sibling progeny from *rict-1(-)* mothers carrying (*oyEx713[ges-1p::rict-1(+)]*), an array that expresses *rict-1(+)* in the intestine. Black circles indicate the means across independent replicate experiments (these replicate experiments are shown in different colors), while the smallest circles indicate gonad arms (n = number of gonad arms scored), also summarized below each violin. The violin displays the overall distribution of data points. Error bars indicate the standard error of the mean. Statistical results and p-values are summarized in Tables S6 and S7, respectively. The asterisks indicate *** p<0.001, ** p<0.01, * p<0.05 of a Linear Mixed Effects Model with a Tukey Post Hoc Test.

To determine whether intestinal *rict-1(+)* is sufficient to promote accumulation of the larval PZ pool, we expressed *rict-1(+)* from an intestinal promoter, *ges-1p,* on an extrachromosomal array in the *rict-1(-)* background. We found that the array-bearing siblings contain a significantly greater number of PZ nuclei (145.2 ± 3.73) than their non-array-bearing siblings (100.6 ± 3.48). In addition, there is no difference between the PZ counts of non-array bearing progeny of array-bearing mothers and homozygous *rict-1(-)* mutants (Tukey comparison p=0.533). This result supports the conclusion that maternal effects do not contribute to *rict-1* regulation of the PZ (Fig. 4B).

Taken together, our necessity and sufficiency tests demonstrate that intestinal RICT-1 is a major contributor to the accumulation of the larval PZ pool. Interestingly, intestinal degradation of RICT-1 does not lower the number of PZ nuclei to the level of the *rict-1(-)* (Fig. 4A), nor does intestinally-expressed *rict-1* entirely rescue to wild-type levels (Fig. 4B). These results indicate that either degradation and heterologous rescue do not reach null or wild-type levels, respectively, or that RICT-1 acts in additional tissues.

Single cell RNA-seq analysis (Fig. S3A, data from (Cao et al., 2017; Hutter and Suh, 2016)) indicates that in addition to the intestine, larval *rict-1* is expressed in other tissues, including the germ line. Given our prior results indicating a role for TOR in the germ line (Korta et al., 2012) we tested RICT-1 germline degradation using *ieSi38*[*sun-1p*::TIR1] and found that it also contributes, though less than the intestine (Fig. S3C). We conclude that although other tissues contribute, the intestine is the primary source of RICT-1 that is important for establishment of the early adult germline PZ, with additional contribution from the germ line.

### Transcripts of four vitellogenin-encoding genes are depleted in the *rict-1(-)* mutant relative to wild type at the mid-L4

To determine how *rict-1* influences the larval germ line, we performed comparative bulk RNA-sequencing in biological triplicate of three strains of tightly synchronized mid-L4 stage worms: wild type, the *rict-1(-)* mutant, and the *rict-1(-)* mutant in which *rict-1(+)* is expressed solely in the intestine. We integrated the extrachromosomal array and determined that the resulting strain was functional and displayed a similar PZ phenotype to worms with *Ex[intestinal rict-1(+)]* (Fig.S3C). This combination of strains allowed us to explore the impact of both global and intestine-specific activity of *rict-1* on the transcriptome. The PCA analysis for all three strains indicates that all three strains are distinctly different from each other (Fig. 5A).

**Figure 5:**
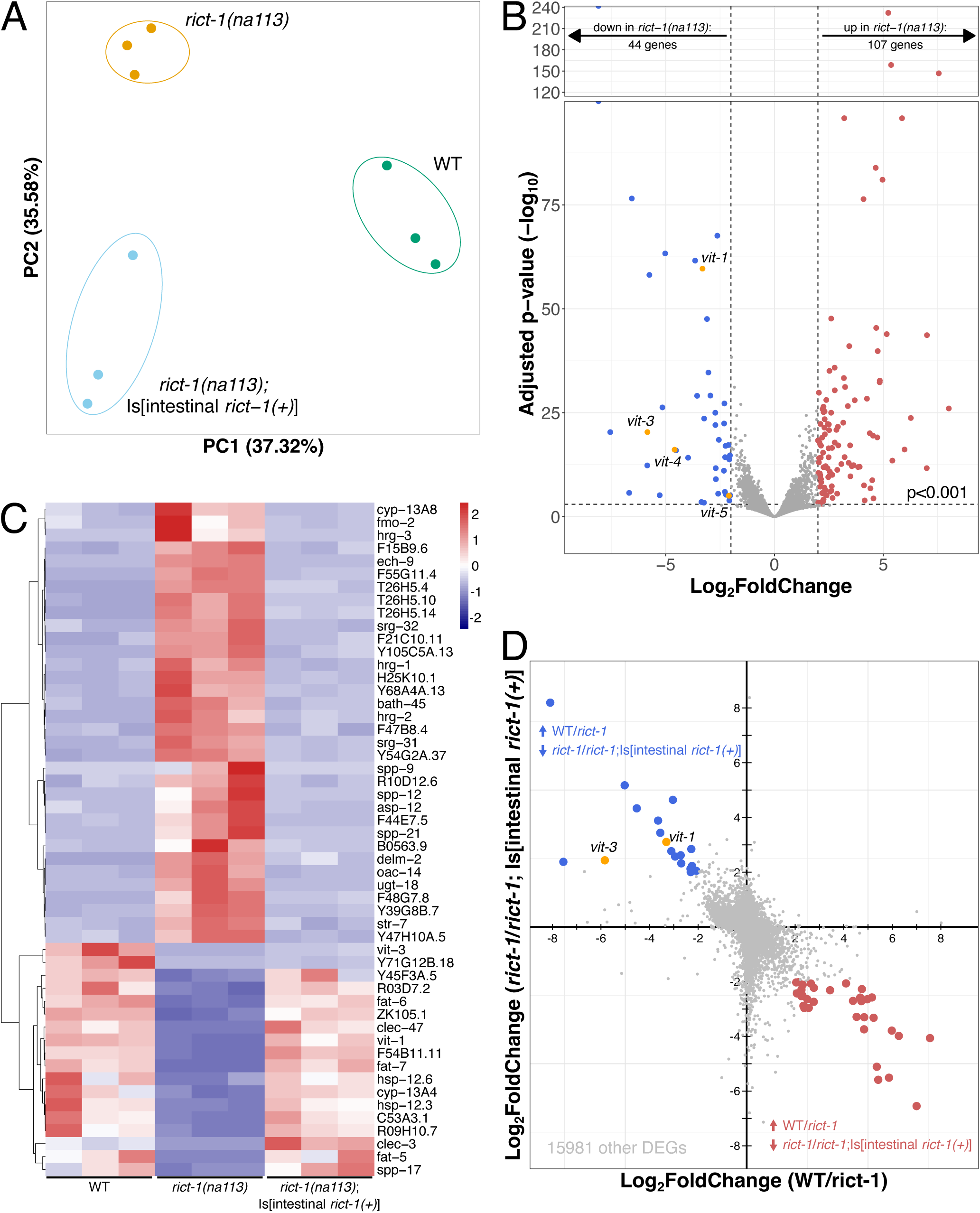
RNA-sequencing reveals that two vitellogenin encoding genes are regulated by intestinal RICT-1. (A) Principal Component Analysis of the three strains, in which each filled circle indicates one library, color coded by the three different strains. (B) Volcano plot displaying a comparison of transcript abundance between *rict-1(-)* mutants and wild type (WT) as circles in both the *rict-1(-)* mutant and the WT: not significantly regulated (grey), down-regulated in the *rict-1(-)* mutant (blue – 107 genes) or up-regulated in the *rict-1(-)* mutant (red – 44 genes) relative to WT. Vitellogenin genes are labeled and marked in orange circles. The horizontal dashed line indicates the adjusted p-value threshold of <0.001, while the vertical dashed lines indicate a log2 Fold Change of −2 or 2. (C) Heatmap in which each line indicates one differentially regulated gene (log2 fold change >2 and adjusted p value < 0.001 – colored points from panel B) between WT, the *rict-1(-),* and the *rict-1(-)* mutant with *Is[intestinal rict-1(+)].* The color scale indicates the Fragments per Kilobase of transcript per Million mapped reads (FPKM), where darker colors indicate a stronger differential, where blue colors indicate lower and red higher abundance. (D) The fold change between WT and *rict-1(-)* is displayed on the x-axis and fold change between *rict-1(-)* and *rict-1(-); Is[intestinal rict-1(+)]* are shown. Blue circles indicate those transcripts of < −2 log2 Fold Change between WT and *rict-1(-)* and >2 Fold Change between *rict-1(-)* and *rict-1(-); Is[intestinal rict-1(+))].* Red circles indicate those transcripts of >2 log2 Fold Change between WT and *rict-1(-)* and < −2 Fold Change between *rict-1(-)* and *rict-1(-); Is[intestinal rict-1(+))]. vit-1* and *vit-3* are labelled in orange circles. Grey circles indicate those transcripts that did not fall into the above-mentioned categories (15981 transcripts). All transcripts can be found in Table S4.

Comparing the *rict-1* mutant and the wild type, we found that among 16,034 protein coding transcripts detected (see Methods), 151 transcripts were differentially abundant (adjusted p < 0.05 and log2 fold change >2), with 44 less abundant and 107 more abundant in the *rict-1(-)* mutant than in the wild type (Fig. 5B). Among the genes with lower transcript abundance in the mutant, the vitellogenin genes caught our attention for several reasons. The *C. elegans* genome encodes six vitellogenins, members of a family of conserved yolk proteins (reviewed in (Perez and Lehner, 2019)) that are best described for their adult function provisioning oocytes (Grant and Hirsh, 1999; Kimble and Sharrock, 1983). However, no function in larval stages or the larval germ line is yet described. We found that the steady-state mRNA levels of four of the six vitellogenins (*vit-1, −3, −4,* and -*5*) are significantly lower in the mid-L4 *rict-1* mutant relative to wild type (adjusted p<0.001, log2 fold change < −2) (Fig. 5B, orange circles).

To assess the contribution of intestinal *rict-1(+)*, we then analyzed the 151 transcripts to determine which of these were significantly different between the mutant alone and the strain expressing *rict-1(+)* solely in the intestine (Fig. 5C). We found that a third of these 151 transcripts were significantly different between the global *rict-1* mutant versus *rict-1* mutants expressing *rict-1(+)* only in the intestine: 18 were higher and 35 were lower for *rict-1(-)* versus *rict-1(-); Is[intestinal rict-1(+)]* (Fig. 5D, upper left and lower right, respectively). Interestingly, transcripts of two of the *vit* genes (*vit-1* and *vit-3)* are significantly higher in the *rict-1(-)* mutant expressing intestinal *rict-1(+)* relative to the *rict-1(-)* mutant alone (Fig. 5D), suggesting that their expression may be regulated by intestinal *rict-1*.

### VIT-1 and VIT-3 promote germline progenitor pool establishment

We wished to determine the functional relevance of VIT-1 and VIT-3 to progenitor zone establishment in larvae. Among the vitellogenin genes we detected in our comparative RNA-seq analysis, *rict-1* was previously found to indirectly regulate *vit-3* transcription in the adult intestine (Breen et al., 2024; Dowen et al., 2016). However, no previous function has been described in larvae for any of the vitellogenins, and no previous connection has been made between *vit-1* and *rict-1*.

We therefore tested whether VIT-1 and VIT-3 could, like RICT-1, contribute to PZ pool establishment, by counting the number of PZ nuclei in worms bearing loss-of-function alleles of *vit-1* or *vit-3*. We observed that *vit-1(-)* and *vit-3(-)* null mutants each contain significantly fewer PZ nuclei (138.1 ± 4.84 and 130.45 ± 2.30, respectively) than the wild type (179.6 ± 5.62 and 202.9 ± 5.35) (Fig. 6A,B).

**Figure 6:**
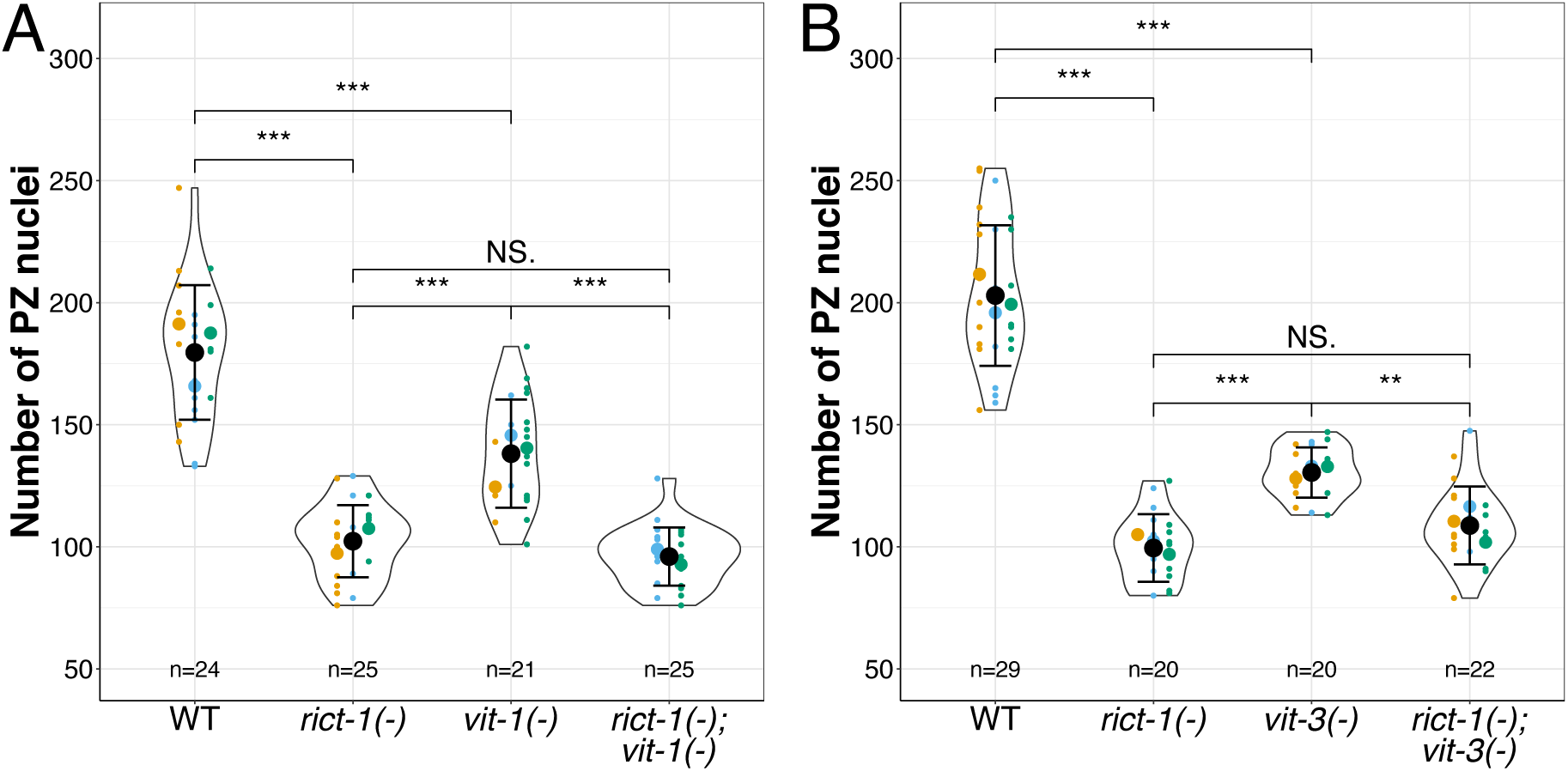
Two vitellogenins, VIT-1 and VIT-3, are likely acting in a linear pathway with RICT-1 to promote establishment of the PZ pool. (A, B) Numbers of PZ nuclei in the wild type (WT) and indicated genotypes. Black circles indicate the means across independent replicate experiments (these replicate experiments are shown in different colors), while the smallest circles indicate gonad arms (n = number of gonad arms scored), also summarized below each violin. The violin displays the overall distribution of data points. Error bars indicate the standard error of the mean. Statistical results and p-values are summarized in Tables S6 and S7, respectively. The asterisks indicate *** p<0.001, ** p<0.01, * p<0.05 of a Generalized Linear Model with a Tukey Post Hoc Test.

If *rict-1* and either *vit* gene were acting in a linear genetic pathway, we would expect that loss of both *rict-1* and either *vit* gene would not further deplete the PZ pool beyond either the *rict-1* or *vit* single mutant alone. We found that the double mutant *rict-1(-); vit-1(-)* and the double mutant *rict-1(-); vit-3(-)* each display the same number of PZ nuclei as the *rict-1(-)* mutant alone (96.0 ± 2.38 for *rict-1(-); vit-1(-) versus* 102.3± 2.96 for *rict-1(-)* alone, and 108.7 ± 3.40 for *rict-1(-); vit-3(-)* versus 99.5± 3.10 for *rict-1(-)*). Together, these results are consistent with a linear genetic pathway and reveal an unexpected role for *vit-1* and vit*-3* in the larval accumulation of germline progenitors, likely downstream of *rict-1*.

## DISCUSSION

This work establishes a developmental context for a novel non-autonomous activity of intestinal RICT1/Rictor on a stem cell system. In addition, it indicates a role for vitellogenins downstream of intestinal RICT-1 in establishment of the germline progenitor pool. In the course of this work, we generated seven new alleles of *rict-1*, including six similarly-behaving alleles encoding small deletions or early stop codons, and one encoding a functional AID-tagged RICT-1 protein. Using these and prior existing alleles, we show that RICT-1 is required for several aspects of germline development and reproduction.

Using multiple mutant alleles of *rict-1*, we confirmed low broods reported by others (Jones et al., 2009; Soukas et al., 2009), and we also confirmed an extended period of progeny production (Webster et al., 2013). Interestingly, we observed no increase of progeny production upon replete sperm provision, suggesting that sperm is not the limiting factor for offspring production in the *rict-1* mutant. In mated worms, brood size correlates with an increased number of cells in the PZ pool, suggesting that the number of PZ cells destined to become oocytes is limiting (Agarwal et al., 2018; Kocsisova et al., 2019). We speculate that a similar limitation could account for our observation.

We focused on a larval requirement for RICT-1 in the accumulation of germline progenitors and determined that the *rict-1* mutant PZ pool reaches half the normal number of cells at the adult molt. Multiple *rict-1* mutants affecting different areas of the protein displayed similarly reduced PZ numbers (∼half of the wild-type). The N-terminus and third exon mutants disrupt only isoform A and B but nevertheless display the same PZ phenotype as those mutants that affect all three isoforms. This suggests that the C isoform is unlikely to play a specific role in the establishment of the PZ.

We observed a similarly-reduced PZ phenotype upon loss of SINH-1, a part of the canonical TORC2 complex, and SGK-1, a canonical TORC2 effector, and found evidence for genetically linear relationships with *rict-1.* By contrast, we find no obvious genetic interaction with the DAF-2/IIS or DAF-7/TGF-ß pathways. We find that RICT-1 promotes robust larval, but not adult, germline cell cycle progression but does not affect the stem cell fate decision.

While both TORC1 and TORC2 are implicated in progenitor pool establishment, our work reveals interesting differences. We previously found that loss of RSKS-1/S6K, a presumed downstream effector of TORC1, has a similarly reduced PZ as RICT-1, and that the requirement for *rsks-1(+)* is germline-autonomous (Korta et al., 2012). By contrast, although we observed a minor germline-autonomous effect, *rict-1* has a strong non-autonomous (intestine-specific) effect on the germ line. In addition, loss of either *rict-1* or *rsks-1* results in reduced larval, but not adult, mitotic index. The reduced larval mitotic index was observed at a time when the cell cycle is relatively short and the mitotic index higher, while the unchanged adult mitotic index was observed at a time when the cell cycle is longer and the mitotic index lower (Roy et al., 2016). However, in contrast with *rsks-1*, loss of *rict-1* does not affect the stem cell fate decision. That is, while loss of *rict-1* additively reduces the number of PZ nuclei in combination with *glp-1(ts)* at a semi permissive temperature, unlike *rsks-1,* it does not enhance the “Glp-1-like” phenotype such that the entire PZ is lost to differentiation. We conclude that RICT-1 promotes robust larval cell cycle progression but not the stem cell fate decision. We nevertheless found that this larval cell cycle role impacts the borders of the early adult stem and non-stem progenitor pools.

We discovered a prominent non-autonomous role for intestinal *rict-1* in larval establishment of the germline PZ that is both necessary and sufficient for establishment of the full wild-type PZ pool. The necessity experiments were conducted using an auxin-mediated protein degradation system. Because auxin exposure prolongs lifespan (Loose and Ghazi, 2021) and promotes resistance to endoplasmic reticulum stress (Bhoi et al., 2021), we used a minimal concentration of auxin (1mM) that did not enhance the PZ defect in the *rict-1* mutant nor in other control strains. RICT-1 intestinal sufficiency has been demonstrated for other phenotypes including entry to the dauer stage, foraging behavior (O’Donnell et al., 2018), associative learning (Sakai et al., 2017), fat accumulation (Jones et al., 2009; Soukas et al., 2009), and food seeking (McLachlan et al., 2022). Our results indicate that the intestinal rescue of *rict-1(+)* in the *rict-1(-)* background does not rescue the number of PZ nuclei to the full wild-type level, and that the intestinal degradation does also not reduce the number PZ nuclei from the wild-type level to the *rict-1(-)* level, a difference that is likely attributable to the additional germline-autonomous role for *rict-1* (Fig. S2C). Whether the intestine and germ line are the only required tissues and how the germline-autonomous role interacts with TORC1 function are areas for further investigation.

Our comparative RNA-sequencing analysis of L4 larvae identified four vitellogenin genes among transcripts that are less abundant in *rict-1* relative to wild type, among which *vit-1* and *vit-3* are partially restored by expression of intestinal *rict-1(+)* in the *rict-1(-)* mutant background. Published RNA-seq analyses of *rict-1* day 1 adults (Breen et al., 2024; VanDerMolen et al., 2025) found significant differential expression of *vit-1*, *vit-2*, *vit-3*, and *vit-4*. RICT-1 indirectly promotes high levels of *vit-3* transcription in the adult intestine (Breen et al., 2024; Dowen et al., 2016).

Although no function has been previously ascribed to vitellogenins in larval stages nor in the germ line prior to adult oocyte provisioning, many studies have examined *vit* regulation, including extensive promoter dissection of *vit-2* (reviewed in (Perez and Lehner, 2019)). In addition, in the L4, *vit-3* expression is regulated by *ins-37*, while *wrt-6* modestly regulates both *vit-1* and *vit-3* (Abete-Luzi et al., 2020). In general, expression of all vitellogenins is low to negligible during early larval stages and increases rapidly during the fourth larval stage and into adulthood, but individual *vit* gene expression dynamics varies somewhat between each gene and across life stages (Boeck et al., 2016; Cao et al., 2017; Heine and Blumenthal, 1986; Hillier et al., 2009; Hutter and Suh, 2016). Taken together, these studies suggest the possibility that despite high sequence similarity between the six vitellogenin-encoding genes (Fig. S4), the *vit* genes may have additional and differential functional roles during larval development.

We demonstrate a functional role for *vit-1* and *vit-3* in promoting progenitor pool establishment, likely downstream of *rict-1.* The *vit-1* and *vit-3* mutant PZ defects are similar to, though not as severe as, loss of *rict-1,* and loss of either *vit* mutant does not further enhance loss of *rict-1*. Although loss of either *vit-1* or *vit-3* causes a mutant phenotype somewhat less severe than *rict-1*, other *vits* identified in our RNA-seq analysis, or other *rict-1*-regulated gene products may contribute to PZ expansion. In support of the former hypothesis, we note that the rapid increase in all *vit* gene expression during the L4 stage correlates with the time of rapid expansion of the PZ pool (Killian and Hubbard, 2004).

We speculate that RICT-1 regulates the establishment of the PZ via intestine-germline transport of lipids or other molecules associated with vitellogenin-containing lipid particles in larval stages. While molecules associated with oocyte-bound vitellogenins likely act as a nutrient source for post-embryonic survival (reviewed in (Perez and Lehner, 2019)), our results suggest that vitellogenins may promote germ cell cycle in earlier stages. One possibility is that vitellogenins facilitate transport of critical nutrient lipids, such as cholesterol. Another possibility is that they provide limiting membrane lipids required for the demands of germ cell division. Yet another possibility is that they associate with as-yet unknown moieties such as hormones or with microRNAs which are part of yolk (Aupérin et al., 2025) to direct germline development. The oocyte receptor for vitellogenins, RME-2, is not detectable by immunohistochemistry in the distal gonad during larval stages (Grant and Hirsh, 1999; Salinas et al., 2007), though it is detected in larval stages by both bulk (Boeck et al., 2016; Hillier et al., 2009; Hutter and Suh, 2016) and single-cell RNA-seq (Cao et al., 2017; Hutter and Suh, 2016) studies. However, a function for RME-2 prior to oogenesis has not been tested. Alternatively, a different receptor may fulfill this role in larvae. Additional studies are required to determine how larval vitellogenins participate in the establishment of the PZ pool.

In conclusion, we present the first evidence that TORC2 contributes to establishment of the full germline stem/progenitor pool. Our results suggest a model whereby intestinal RICT-1 promotes the expression of vitellogenins that support germline progenitor pool expansion during larval stages. Due to the high conservation of TORC2 components, vitellogenins, and their receptors, our results may have general implications for how Rictor and TORC2 regulate germline development in other organisms and, more generally, how they may support pools of proliferating stem or progenitor cells in other organisms and organ contexts. Moreover, since vitellogenin-like molecules, such as ApoB, are implicated in heart disease and other aspects of lipid metabolism (reviewed in (Sniderman et al., 2019)), the *C. elegans* larval germ line may provide inroads to the identification of additional associated moieties and receptors.

## ACKNOWLEDGMENTS

We thank the Microscopy (RRID: SCR_017934) and Genome Technology (RRID: SCR_017929) cores at NYU Grossman School of Medicine. The Microscopy Core is partially supported by the Cancer Center Support Grant P30CA016087 and P01A1080192 at the Laura and Isaac Perlmutter Cancer Center. The Genome Technology Core is partially supported by the Cancer Center Support Grant P30CA016087 at the Laura and Isaac Perlmutter Cancer Center. We thank Nicole Hall for help and advice with the RNA extraction protocol and analysis, Amy Webster for feedback on the RNA-seq analysis, Sophie Wasel and Mia Sinks for technical support, and Michael Cammer for microscopy support. We are also grateful for feedback and support from Jeremy Nance and Niels Ringstad, and members of their labs. The computational requirements for this work were supported in part by the NYU Langone High Performance Computing (HPC) Core’s resources and personnel. Thanks to Michael O’Donnell for providing strains; some strains were also provided by the CGC, which is funded by NIH Office of Research Infrastructure Programs (P40 OD010440). This work was supported by NYSTEM Institutional Training Grant Contract #C322560GG and NIH R35GM134876 to E.J.A.H., and a postdoctoral fellowship by the American Heart Association to AK (https://doi.org/10.58275/AHA.25POST1362264.pc.gr.227408).

## Competing Interests

The authors declare no competing interests.

## MATERIALS AND METHODS

### Worm maintenance and husbandry

*C. elegans* nematodes were grown on NGM solid media at 20°C and fed *Escherichia coli* strain OP50 (Stiernagle, 2006). All strains generated and used in this study are listed with full genotypes in Table S1, including the oligonucleotides used for strain validation. The sequences of these oligonucleotides are listed in Table S2. Some strains were provided by the CGC. All strains were based on Bristol N2 (RRID:WBSTRAIN: WBStrain00000001). Many experiments required access to information on WormBase (Sternberg et al., 2024). The following strains were generated in this study: GC1745, GC1746, GC1747, GC1755, GC1756, GC1759, GC1762, GC1764, GC1779, GC1779, GC1780, GC1793, GC1801, GC1802, GC1815, GC1816, GC1829, GC1832, GC1842, GC1886, GC1893, GC1905.

### Determination of progenitor zone nuclei counts and mitotic index

Mid-L4 worms (L4.5; (Mok et al., 2015)) were picked from mixed stage plates and transferred to new plates. 6h later all early adult worms bearing a vulval slit were picked into 1ml of M9+Tween within a 1.5ml of low-retention microfuge tube (02-681-320, Fisherbrand). Worms were spun down on a table-top centrifuge for 10 seconds, before the supernatant was carefully removed with a pipette under the dissecting scope, to avoid loss of worms. Another 500µl of M9+Tween was added, followed by a centrifugation and supernatant removal. 50µl of 100%EtOH were added to each sample, and worms were allowed to settle at the bottom for ∼10min. After 10min as much EtOH was removed as possible without disturbing the worms, and ∼5µl of Vectashield+DAPI (H-1000, Vector Laboratories) was added. Worms were gently pipetted onto a 4% agar pad on a glass slide. Slides were kept in the cold and dark and were imaged within 2 weeks after staining. Imaging was performed on Nikon W1 spinning disk confocal microscope with 60x oil immersion magnification with z-stacks of 0.5µm step size. Only one gonad arm was counted per individual worm. PZ nuclei were manually counted using the point tool in ImageJ (Arganda-Carreras et al., 2024). The PZ border was determined as the distal-most row with 2 or more crescent-shaped nuclei typical of the transition zone (Crittenden et al., 2006). Mitotic index is presented as a percentage of the number of mitotic figures (characteristic metaphase and anaphase figures) divided by the total number of PZ nuclei.

### Brood Size Determination

L4 worms were picked from mixed stage plates. Single hermaphrodites were added to plates and, for the mated treatment, were joined by 3 wild-type males, which were replaced every two days. In mating experiments (Fig. S1), mating was confirmed by abundant male offspring (N2 males) and/or mating plug (CB4855 males). Mated *rict-1(-)* hermaphrodites exhibited poorer survival than self-fertile hermaphrodites; however, offspring counts are included only from worms that lived several days after thelast progeny were produced (see Fig. S1). All worms were transferred every day to new plates, and hatched offspring were counted 2 days later.

### CRISPR-Cas9 genome editing

CRISPR/Cas9 genome editing was performed using pre-incubated Cas9 (Berkeley)::(crRNA +tracrRNA) (IDT) ribonucleoprotein, and screening was facilitated by co-CRISPR detection using the *dpy-10(cn64)* mutation as described (Paix et al., 2017). DNA repair templates contained ∼25–35 bps of homology on each arm as ssDNA oligos (<150 bps) (IDT). The guide RNAs and oligos are listed in Table S3. For Stop-In mutants (*na112, na113, na114*, and *na115*) the inserted stop cassette led to an early stop in all frames (Wang et al., 2018), while the sAID tag did not interfere with the function of RICT-1 (Fig. S3B).

### Integration of extrachromosomal array

The extrachromosomal array *oyEx713* in GC1756 was integrated using x-rays, as previously described (Evans, 2006).

### RNA extraction for bulk RNA-seq

Worms were bleached and synchronized as previously described (Stiernagle, 2006). Due to differences in growth rate (as previously empirically determined), synchronized L1 larvae were added to 15 10cm NGM plates (each inoculated with 1ml of OP50) at different times so that all strains would reach the mid-L4 stage at the same time. Worms were washed off the plates with RNAse-free water (10977015, Invitrogen) and collected atop pluristrainers with 40µl mesh size (#43-10040-40 from Pluristrainer.com) and washed three times with 5ml of RNAse free H_2_0. Pluristrainers were turned over, and 1ml of RNAse-free water was used to wash the worms off the mesh and collect them in RNAse-free 2ml Eppendorf microfuge tubes (E0030123620, Eppendorf). Each 1ml of packed worms was resuspended in 4ml of Trizol. The worm+Trizol mix was vortexed and flash-frozen by carefully dropping into liquid nitrogen and then thawed at 37°C. After three freeze-thaw cycles, samples were thawed at 37°C, and 2ml of Trizol+2ml of chloroform per packed ml of worms was added and vigorously shaken by hand. Samples were aliquoted into 2ml RNAse-free tubes and incubated at room temperature for three minutes. Samples were then spun at 12,000g at 4°C for 15 min. The aqueous layer was removed into new RNAse-free tubes, and an equal volume of 70% ethanol was added and inverted to mix. This solution was transferred to a spin column (RNeasy Qiagen kit # 74104) and was spun down at 21k rcf for 30sec. The column was washed once with buffer RW1 and centrifuged for 15 seconds at 21k rcf DNase I was added directly to each RNeasy column membrane and was left to incubate at room temperature for 15 minutes. Each column was washed with Qiagen RW1 and RPE buffers, before the sample was eluted from the spin column in RNase-free water to be collected in a fresh 2ml RNase-free tube. RNA quality was initially assessed with a nanodrop (NanoDrop 8000 UV-Vis Spectrophotometer). Samples were submitted to the NYU genome technology core where RNA extractions were quantified using RNA Nano Chips (Cat. #5067-1511) on an Agilent 2100 BioAnalyzer. RNA-Seq library preps were constructed with the Illumina TruSeq Stranded mRNA Library Prep kit (Cat #20020595) according to manufacturer’s recommendations. The input amount of total RNA was 450ng with 12 cycles of amplification. Final libraries were visualized using High Sensitivity DNA ScreenTape (Agilent, Cat. #5067-5584) on the Agilent TapeStation 2200 instrument. Quant-It (Invitrogen, Cat. P11495) was used for concentration determination and libraries were pooled equimolar. The pool was sequenced paired-end 50 cycles on an Illumina NovaSeq X+ 10B 100 cycle flowcell with 2% PhiX spike-in.

### RNA-seq analysis

Version WBCel235 of the *C. elegans* genome was used to map sequencing reads (Table S4). Seq-N-Slide (Dolgalev, 2024) was used to align the sequencing results and determine those gene transcripts that are differentially expressed between the mutant and wild-type strains. Transcript counts were restricted to protein coding genes (19,983 genes). Mapping Efficiency is provided in Table S5. Prior to further analysis this count data was further restricted to include only genes represented in at least 2/3 libraries per strain with non-zero counts (16,034 genes). Principal component analysis (PCA) was performed on log2 normalized CPM values on all genes included in the differential gene expression analysis (16,034 genes), using the stats package in R (version 2024.04.2+764) (RStudio Team, 2020). Lists of genes with differentially abundant transcripts between WT versus *rict-1* and *rict-1* versus *rict-1; Is[intestinal rict-1(+)]* can be found in Table S4. The entire dataset will be publicly available at NCBI Gene Expression Omnibus.

### Statistical analyses

Plots in Figures 1-6 and Figures S1-S3 were generated using R with minor modifications using Adobe Illustrator or Inkscape. Statistical analysis was conducted in R (version 2024.04.2+764) (RStudio Team, 2020). All PZ counts and the offspring data were analyzed with a Generalized Mixed Effect Model (Nelder and Wedderburn, 1972), taking the replicate as a random factor into account. The Mitotic Index data was analyzed using a Linear Model (Nelder and Wedderburn, 1972), and the Reproductive Period data was analyzed with a Linear Mixed Effects Model (Nelder and Wedderburn, 1972). All results from the statistical tests can be found in Table S6 and all detailed p-values can be found in Table S7. Summaries of all data are in Table S8.

Conceptualization: A.K., E.J.A.H.;

Formal analysis: A.K.

Funding acquisition: A.K., E.J.A.H.;

Investigation: A.K.

Methodology: A.K., E.J.A.H.;

Project administration: E.J.A.H.;

Supervision: E.J.A.H.;

Visualization: A.K.

Writing original draft: A.K.

Writing – review & editing: A.K., E.J.A.H.;

## Notes

### Competing Interest Statement

The authors have declared no competing interest.

### Summary of Updates

Additional data and data analysis are included in this revised version affecting all 6 main figures. Supplemental information is also updated.

