## Supplemental material for "Intestinal RICT-1/Rictor regulates larval germline progenitors via vitellogenins VIT-1 and VIT-3 in *C. elegans*"

### SUPPLEMENTAL FIGURES

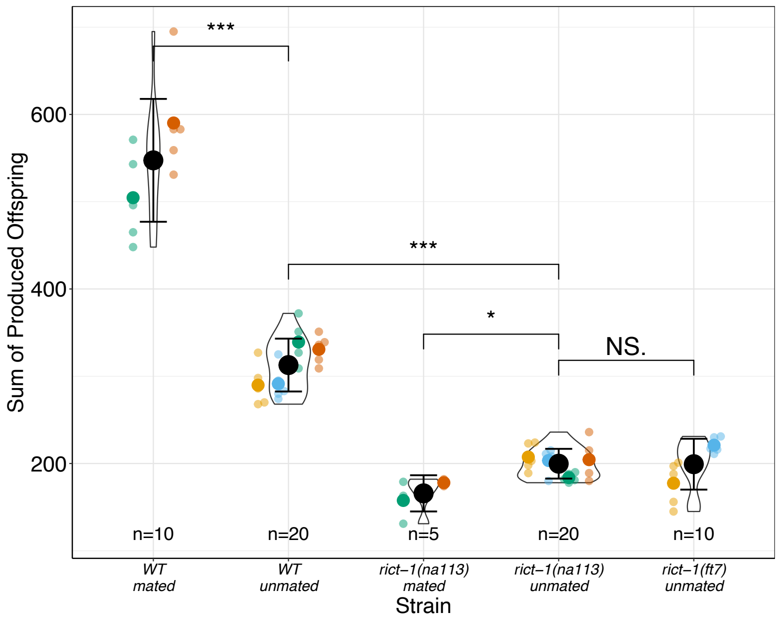


Figure S1: Mated *rict-1* mutant hermaphrodites do not produce more offspring than self-fertile hermaphrodites. Numbers of hatched offspring from unmated hermaphrodites and mated hermaphrodites. The data for unmated worms here are the same as those displayed in Fig. 1E-G. Worms were mated with N2 (in orange) or CB4855 (in red) males. Black circles indicate the means across different replicate experiments (shown in different colors), while the smallest circles indicate individual gonad arms (n = number of gonad arms scored). The violin indicates the overall distribution of data points. Error bars indicate the standard error of the mean. Two-sided t-tests were performed, and the significances are noted as follows: *** $p<0.001$, * $p<0.05$.

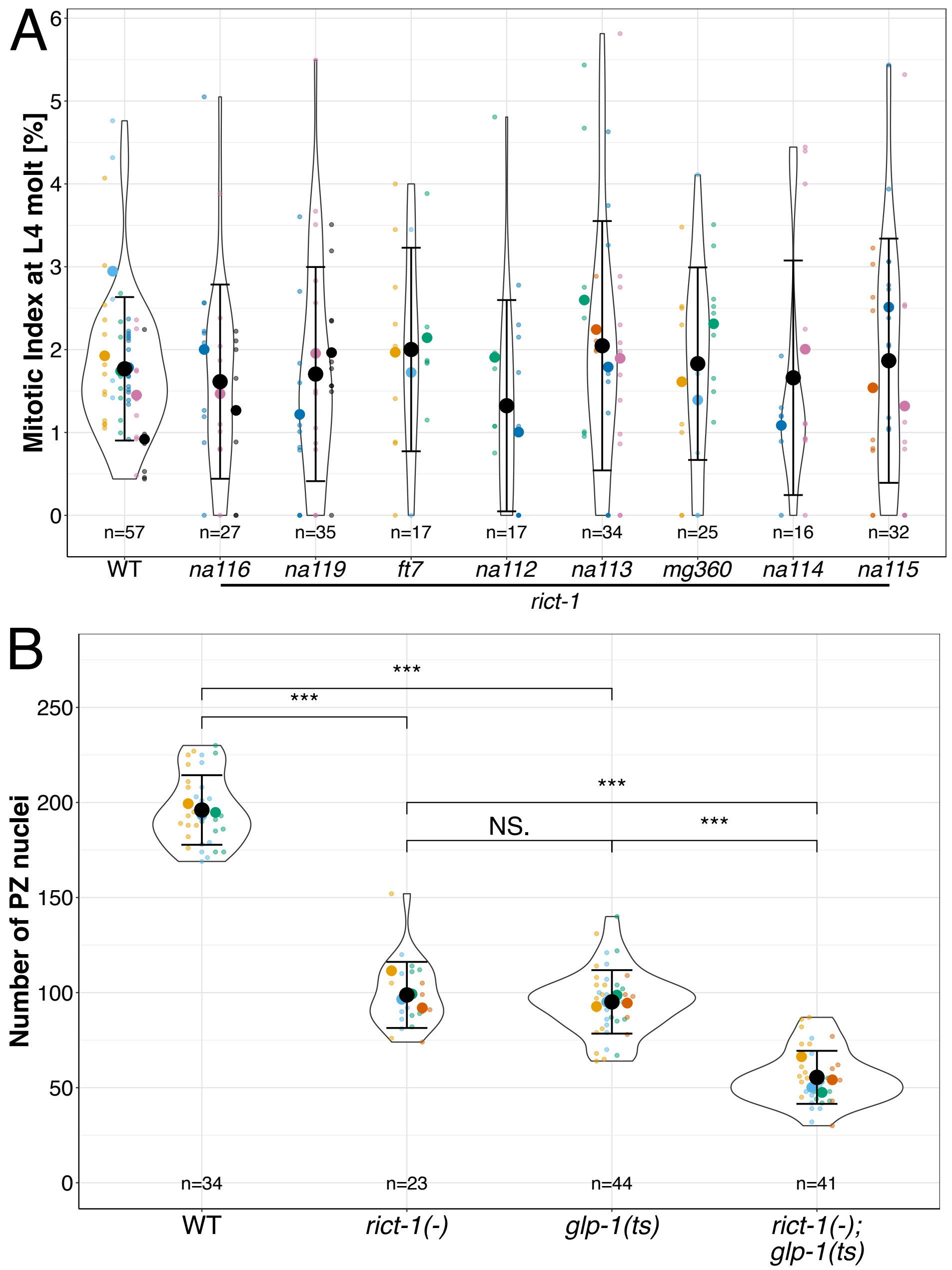


Figure S2: RICT-1 does not affect the mitotic index at the L4 to Adult molt and likely acts in parallel to GLP-1. (A) Mitotic index in adult wild type (WT) and indicated *rict-1* mutants. (B) Numbers of PZ nuclei in wild type (WT), *rict-1(na113)*, *glp-1(e2141)*, and the double mutant *rict-1(na113); glp-1(e2141)* at the L4 to Adult molt. Black circles indicate the means across independent replicate experiments (shown in different colors), while the smallest circles indicate individual gonad arms (n = number of gonad arms scored). The violin indicates the overall distribution of data points. Error bars indicate the standard error of the mean. Statistical tests are summarized in Table S2. The asterisks indicate *** p<0.001 of a Linear Mixed Effects Model with a Tukey Post Hoc Test.

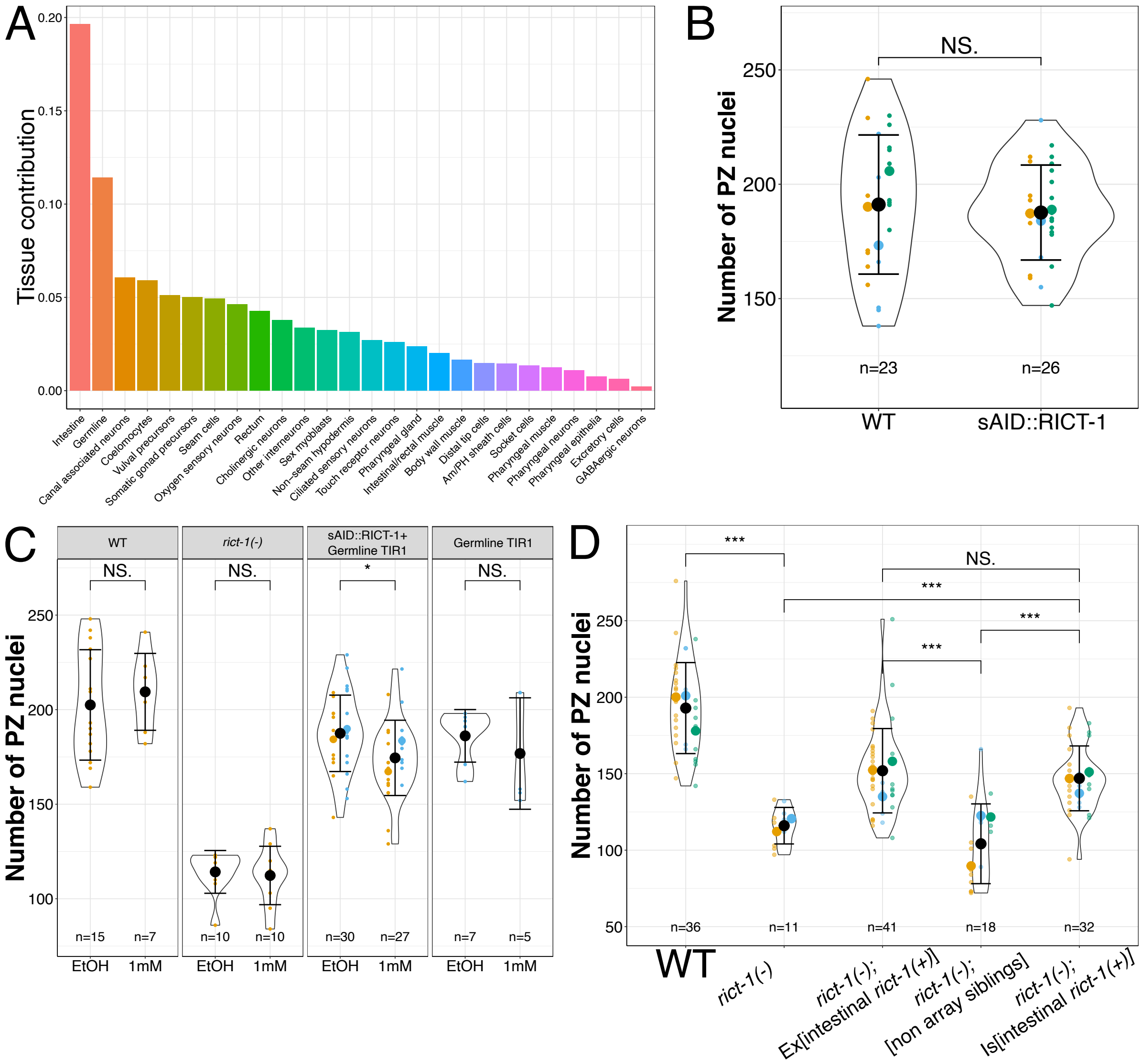


Figure S3: RICT-1 is expressed in many tissues, the *na117* [sAID::RICT-1] fusion does not affect the PZ phenotype, RICT-1 is acting on the PZ from the germline albeit at a smaller effect than from the intestine, and the *rict-1(na113)* mutant with an integrated intestinal *rict-1(+)* used in the RNA-seq experiment does not affect the number of PZ nuclei. (A) Proportion of *rict-1* transcripts as a function of tissue assignment from L2 scRNA-seq (Cao et al., 2017; Hutter and Suh, 2016). The relative contribution of each tissue is shown. (B) Number of PZ nuclei in wild type (WT) and GC1802 *rict-1(na117*[sAID::RICT-1]). (C) Number of PZ nuclei in wild type (WT) and strains that deplete RICT-1. Germline TIR1 is *ieSi38* [*sun-1p*::TIR1]. (D) Numbers of PZ nuclei in WT, *rict-1*(*na113*) mutants, array bearing and non-array bearing siblings from mothers carrying *oyEx713[ges-1p::RICT-1 cDNA]*, and a strain generated from the integration of the same array. Data for all other strains are the same as in Fig. 4B. (B, C, D) Black circles indicate the means across different replicate experiments (shown in different colors), while the smallest circles indicate individual gonad arms (n = number of gonad arms scored). The violin indicates the overall distribution of data points. Error bars indicate the standard error of the mean. The asterisks indicate *** $p<0.001$, * $p<0.05$ of a Generalized Linear Model with a Tukey Post Hoc Test.

| | VIT-1 | VIT-2 | VIT-3 | VIT-4 | VIT-5 | VIT-6 |
|---|---|---|---|---|---|---|
| VIT-1 | 100 | | | | | |
| VIT-2 | 94.05 | 100 | | | | |
| VIT-3 | 56.46 | 57.38 | 100 | | | |
| VIT-4 | 56.71 | 57.63 | 98.69 | 100 | | |
| VIT-5 | 56.84 | 57.7 | 97.88 | 98.13 | 100 | |
| VIT-6 | 31.02 | 31.34 | 31.08 | 31.21 | 31.34 | 100 |

Figure S4: Uniprot comparison of the six vitellogenin protein sequences. Percent of amino acid sequence identity over entire protein sequences (The UniProt Consortium et al., 2025)

### SUPPLEMENTAL TABLES

**Table S1:** Generated worm strains and validation primers used in this study

| Strain | Genotype | Primers used for strain validation (Details in Table S5) | Reference/Source and/or parent strain |
|---|---|---|---|
| CA1199 | *unc-119(ed3)* III; *ieSi38* IV | GCo3175 + GCo3178 | CGC (Zhang et al., 2015) |
| CA1209 | *ieSi61 II; unc-119(ed3)* III | GCo3081 + GCo3202 | CGC (Zhang et al., 2015) |
| CB1376 | *daf-3(e1376)* X | GCo2044 + GCo2045 | CGC (Riddle et al., 1981, 198) |
| CB4855 | *Wild type – cross* | NA | CGC (Hodgkin and Doniach, 1997)11/7/25 4:04:00 PM |
| CF1038 | *daf-16(mu86)* I | GCo3147 + GCo3148 + GCo3149 | CGC (Lin et al., 1997) |
| DLS953 | *rict-1 & pqn-32(rhd314) II* | | Dowen Lab (Breen et al., 2024) |
| GC832 | *glp-1(e2141)* III | GCo1217 + GCo1218 | (Dalfó et al., 2010; Priess et al., 1987); 11/7/25 4:04:00 PM |
| GC1745 | *rict-1(na112)* II | GCo2979 + GCo2980 | This work, CRISPR/Cas9 |
| GC1746 | *rict-1(na113)* II | GCo2979 + GCo2980 | This work, CRISPR/Cas9 |
| GC1747 | *rict-1(na114)* II | GCo2993 + GCo2994 | This work, CRISPR/Cas9 |
| GC1755 | *sygl-1(q983)* I; *rict-1(na113)* II | GCo2979 + GCo2980<br>GCo2941 + GCo2942 | GC1746, JK5893 |
| GC1756 | *rict-1(na113)* II*; oyEx713* [*ges-1p*::*rict-1 cDNA*::SL2::mCherry; *unc-122*p::GFP] | GCo2979 + GCo2980 | GC1746, PY9313 |
| GC1759 | *daf-16(mu86)* I; *rict-1(na113)* II | GCo2979 + GCo2980<br>GCo3147 + GCo3148 + GCo3149 | GC1746, CF1038 |
| GC1762 | *rict-1(na113)* II*; daf-3(e1376)* X | GCo2979 + GCo2980<br>GCo2044 + GCo2045 | GC1746, CB1376 |
| GC1764 | *rict-1(na115)* II | GCo2993 + GCo2994 | This work, CRISPR/Cas9 |
| GC1779 | *rict-1(na113)* II; *sgk-1(ft15)* X | GCo2979 + GCo2980<br>GCo3070 + GCo3071 | GC1746, KQ1564 |
| GC1780 | *rict-1(na113)* II*; sgk-1(ok538)* X | GCo2979 + GCo2980<br>GCo3072 + GCo3073 | GC1746, VC345 |
| GC1793 | *daf-16(mu86)* I; *rict-1(na113)* II; *daf-3(e1376)* X | GCo2979 + GCo2980<br>GCo3147 + GCo3148 + GCo3149<br>GCo2044 + GCo2045 | GC1762, CF1038 |
| GC1801 | *rict-1(na116)* II | GCo3040 + GCo3018 | This work, CRISPR/Cas9 |
| GC1802 | *rict-1(na117)* II [sAID::RICT-1] | GCo3040 + GCo3018 | This work, CRISPR/Cas9 |
| GC1815 | *ieSi61 rict-1(na117)* II; *unc-119(ed3)* III | GCo3040 + GCo3018<br>GCo3081 + GCo3202 | GC1802, CA1209 |
| GC1816 | *rict-1(na117)* II; *unc-119(ed3)* III*; ieSi38* IV | GCo3040 + GCo3018<br>GCo3202 + GCo3275 | GC1802, CA1199 |
| GC1829 | *rict-1(na113)* II; *naIs110* [*ges-1p*::*rict-1 cDNA*::SL2::mCherry; *unc-122*p::GFP] | GCo2979 + GCo2980 | This work; X-ray integration of array. Starting strain GC1756. |
| GC1832 | *rict-1(na119)* II | GCo3040 + GCo2980 | This work, CRISPR |
| GC1842 | *rict-1(na113)* II*; vit-3(ok2348)* X | GCo2979 + GCo2980<br>GCo3223 + GCo3224 | GC1746, RB1815 |
| GC1886 | *sinh-1(pe420) rict-1(na113)* II | GCo3211 + GCo3212<br>GCo2979 + GCo2980 | JN2411, GC1746 |
| GC1893 | *rict-1(na113)* II; *glp-1(e2141)* III | GCo2979 + GCo2980<br>GCo1217 + GCo1218 | GC1746, GC1593 |

| | | | |
|---|---|---|---|
| GC1905 | *rict-1(na113)* II; *vit-1(ok2616)* X | GCo2979 + GCo2980<br>GCo3308+ GCo3309 | GC1746, RB1982 |
| JK5893 | *sygl-1(q983)* I | GCo2941 + GCo2942 | CGC (Shin et al., 2017) |
| JN2411 | *sinh-1(pe420)* II | GCo3211 + GCo3212 | CGC (Sakai et al., 2017) |
| KQ6 | *rict-1(mg360)* II | GCo2993 + GCo2994 | CGC (Jones et al., 2009) |
| KQ1366 | *rict-1(ft7)* II | GCo2979 + GCo2980 | CGC (Jones et al., 2009) |
| KQ1564 | *sgk-1(ft15)* X | GCo3070 + GCo3071 | CGC (Jones et al., 2009) |
| N2 | Wild Type | NA | CGC |
| PY9313 | *rict-1(ft7)* II*; oyEx713*[*ges-1p*::*rict-1 cDNA*::SL2::mCherry; *unc-122*p::GFP] line 2 | GCo2979 + GCo2980 | (O'Donnell et al., 2018) |
| RB1815 | *vit-3(ok2348)* X | GCo3223 + GCo3224 | CGC (The C. elegans Deletion Mutant Consortium, 2012) |
| RB1982 | *vit-1(ok2616)* X | GCo3308+ GCo3309 | CGC (The C. elegans Deletion Mutant Consortium, 2012) |
| VC345 | *sgk-1(ok538)* X | GCo3072 + GCo3073 | CGC (The C. elegans Deletion Mutant Consortium, 2012) |

**Table S2** Sequences of oligonucleotides and CRISPR/Cas9 guide RNAs used for strain validation or generation.

| Name | Sequence |
|---|---|
| GCo1217 | ctcgcaaatggatatcccgg |
| GCo1218 | gccgcagaatacgcagatcc |
| GCo2044 | gccgtttctggtttgacacctgg |
| GCo2045 | cggcaggatccagttgacgatatg |
| GCo2856 | acaacccgcatcaaaacttggtctccaacag |
| GCo2857 | cgctgggaattagattaatcaacggctgacttag |
| GCo2941 | gcataacttccatggagttagag |
| GCo2942 | gttttcgagggtgctgagcacg |
| GCo2979 | TGTTAAAATTCCTACCCGACGATCACCCGT |
| GCo2980 | acttttcagatttattctgcaacgaccaga |
| GCo2993 | ATACTTCTCTGGGAGTTCAGGCGCCATTGC |
| GCo2994 | TCGCATATCGGAACCTATTGGATTCCCATC |
| GCo3018 | CGATTCCACTGTGGTTTCCAAAATCGTC |
| GCo3040 | atattgatttatttcgagttgcttgagaaa |
| GCo3069 | catgaaaaatttcaattttcaggcgATGccaaaagaccctgctaaacctcctgccaaggctcaagttgtaggatggccacctgttagatcatacagaaagaatgtcatggtttcttgtcaaaagtcttcaggtggaGGAGGAGGTGACACTCGTCGAAAAGTGTATCATCGTGGCGAAAACAATACACAGAGGATACAGG |
| GCo3070 | gctacttatttcacgaccaagctgcggaac |
| GCo3071 | tgcaagctgacagcaagttctacga |
| GCo3072 | gacgcgatgaccggaaatatgccagttc |
| GCo3073 | agcttttctctcacctccaacgcgagaa |
| GCo3081 | gctccgatgtgatcctatagtgaatata |
| GCo3147 | ctttctcgcgcctttgtctctctatcgg |
| GCo3148 | GGGACATTTTGCAACAGAGCTCACACAC |
| GCo3149 | gagaggttcgattgagttcggggactga |
| GCo3175 | TCTGCAGAATTCGCCCTTAAATGCAAAAGAGAATCGCCTTGTC |
| GCo3178 | acagatccctcaagaacaaccttcatgcg |
| GCo3202 | tgaagaccttacgacggcac |
| GCo3211 | cccaaacgctgtacatgaagaatctcct |
| GCo3212 | ccaaaatccccgccgaacctcataaata |
| GCo3223 | cggtggaattcctcaacatc |
| GCo3224 | ggatttgcttcaaagaccca |
| GCo3275 | ATTTTTATTACTAAAATCAGTTTCTAAAATG |
| GCo3308 | aaaagtgtcgattcggctctaacgtga |
| GCo3309 | acaacgagcacaggaggtggagaca |
| GCo3334 | AAGCTTTCAAGTTAATTATT |

| | |
|---|---|
| GCo3335 | ATCAAACGGAACGATTGGAAGCTTTCAAGTTAATTTACCCATACGACGTGCCAGACTATGCATACCCATACGACGTGCCAGACTATGCATACCCATACGACGTGCCAGACTATGCAGGGAAGTTTGTCCAGAGCAGAGGTGACTAAGTGATAAGCTAGCATTTGGATGCTTAAAATCTACGAGAAATCCAATCTCAAGAA |
| GCo3336 | TCAAGTATCGGAAGCACAGA |
| GCo3337 | GCGAGCCTGCTGGCAGTATCAAGTATCGGAAGCACTACCCATACGACGTGCCAGACTATGCATACCCATACGACGTGCCAGACTATGCATACCCATACGACGTGCCAGACTATGCAGGGAAGTTTGTCCAGAGCAGAGGTGACTAAGTGATAAGCTAGCAGATGGTGGATTTGAGATCCTTCCCGCTGATGCTGTGCCAG |
| GCo3348 | CGTCGAAAAGTGTATCACA |

**<u>Table S3:</u>** List of guide RNAs and repair oligonucleotides used for CRISPR/Cas9 genome editing. See Table S5 for sequences.

| Plasmid/Oligo | Strain | Method used, result | Injected into | Guide used | Co-CRISPR |
|---|---|---|---|---|---|
| GCo3335 | *GC1745 (rict-1(na112)) GC1746 (rict-1(na113))* | CRISPR, early stop in | N2 | GCo3334 | dpy-10 |
| GCo3337 | *GC1747 (rict-1(na114)) GC1764 (rict-1(na115))* | CRISPR, early stop in | N2 | GCo3336 | dpy-10 |
| NA | GC1801 (*rict-1(na116)*) | CRISPR, deletion | N2 | GCo3348 | dpy-10 |
| GCo3069 | GC1802 (*rict-1(na117)*) | CRISPR, sAID::RICT-1 | N2 | GCo3348 | dpy-10 |
| NA | GC1832 (*rict-1(na119)*) | CRISPR, deletion | N2 | GCo3349 | dpy-10 |

**<u>Table S4:</u>** Transcripts of RNA-seq with the fold change between WT and *rict-1*, and *rict-1* and GC1829 (*rict-1; Is[intestinal rict-1(+)]).* (See Table S4 xlsx file)

Table S5: Efficiency of RNA-seq reads indicated as total reads, uniquely mapped reads for each library.

| Strain | Total input reads | Uniquely mapped reads | Efficiency |
|---|---|---|---|
| WT_1 | 223967331 | 201890297 | 90.14% |
| WT_2 | 165383782 | 151269973 | 91.47% |
| WT_3 | 166932106 | 152483471 | 91.34% |
| *rict-1(na113)*_1 | 225260930 | 203548165 | 90.36% |
| *rict-1(na113)*_2 | 141964671 | 126244017 | 88.93% |
| *rict-1(na113)*_3 | 132369125 | 121846848 | 92.05% |
| *rict-1(na113); Is[intestinal rict-1(+)]*_1 | 181103391 | 158463847 | 87.50% |
| *rict-1(na113); Is[intestinal rict-1(+)]*_2 | 178641027 | 152172742 | 85.18% |
| *rict-1(na113); Is[intestinal rict-1(+)]*_3 | 157314848 | 142438477 | 90.54% |

**<u>Table S6.</u>** Overview of statistical results organized by Figure. The columns indicate the statistical model, the variable, the factor, the random factor, additional info, the Chi-Square, the degrees of freedom, the p-value and the significance. LMEM stands for Linear Mixed Effect Model, GLM for Generalized Linear Model, LM for Linear Model.

| Fig | Statistical Model | Variable | Factor | Random Factor | Additional Info | Chi Sq | Degrees of Freedom | p value | Sig |
|---|---|---|---|---|---|---|---|---|---|
| 1B | LMEM | Number of PZ nuclei | Strain | Replicate | | 1525.1 | 8 | < 2.2e-16 | *** |
| 1E | LMEM | Sum of Produced Offspring | Strain | Replicate | | 247.57 | 2 | < 2.2e-16 | *** |
| 1F | GLM | Hatched offspring over time | Strain | | family= poisson | 591.43 | 2 | < 2.2e-16 | *** |
| 1G | LMEM | Reproductive Period | Strain | Replicate | | 34.916 | 2 | 2.618e-16 | *** |
| 2A | LMEM | Number of PZ nuclei | Strain | Replicate | | 348.89 | 3 | < 2.2e-16 | *** |
| 2B | LMEM | Number of PZ nuclei | Strain | Replicate | | 666.43 | 5 | < 2.2e-16 | *** |
| 3A | LMEM | Number of PZ nuclei | Strain | Replicate | | 726.08 | 6 | < 2.2e-16 | *** |
| 3B | LMEM | Mitotic Index | Strain | Replicate | | 5.01 | 1 | 0.0252 | * |
| 3C | Chi -squared test | Gonad Arm Categories | Strain | | x-squared = 189.83 | | 9 | < 2.2e-16 | *** |
| 3D | T-test | Length of SYGL-1(+) | Strain | | t=5.668, df=166.42 | | | 6.238e-08 | *** |
| 3E | T-test | Length of PZ | Strain | | t=5.664, df=126.54 | | | < 2.2e-16 | *** |
| 3F | T-test | Relative Length of SYGL-1(+) / PZ Length | Strain | | t=-1.1668, df=119.51 | | | 0.2456 | NS |
| 4A | LMEM | Number of PZ nuclei | Strain | Replicate | | 432.82 | 9 | < 2.2e-16 | *** |
| 4B | LMEM | Number of PZ nuclei | Strain | Replicate | | 241.12 | 3 | < 2.2e-16 | *** |
| 6A | LMEM | Number of PZ nuclei | Strain | Replicate | | 273.97 | 3 | < 2.2e-16 | *** |
| 6B | LMEM | Number of PZ nuclei | Strain | Replicate | | 434.05 | 3 | < 2.2e-16 | *** |
| S1 | LMEM | Sum of Produced Offspring | Strain | Replicate | | 772.14 | 4 | < 2.2e-16 | *** |
| S2A | LM | Mitotic Index | Strain | | Sum Sq = 8.29, F value= 0.6 | | 8 | 0.7189 | NS |
| S2B | LMEM | Number of PZ nuclei | Strain | Replicate | | 1422.8 | 3 | < 2.2e-16 | *** |
| S3B | LMEM | Number of PZ nuclei | Strain | Replicate | | 0.4949 | 1 | 0.4818 | NS |
| S3C | LMEM | Number of PZ nuclei | Strain | Replicate | | 225.05 | 7 | < 2.2e-16 | *** |
| S3D | LMEM | Number of PZ nuclei | Strain | Replicate | | 168.55 | 4 | < 2.2e-16 | *** |

**Table S7:** List of comparisons and resulting p-values. (See Table S7 xlsx file)

**Table S8:** Mean, standard deviation, standard error, 95% confidence intervals and n values listed by Figure. (See Table S8 xlsx file)
